# Targeting DNA-PK is a highly conserved poxvirus innate immune evasion mechanism

**DOI:** 10.1101/2024.10.06.616852

**Authors:** Marisa Oliveira, Paul T. Manna, Damaris Ribeiro-Rodrigues, Long Fung Chau, Thomas Allport, Emma Wagner, Heather Brooks, Michael A. Skinner, Efstathios S. Giotis, Amanda K. Chaplin, Rodrigo Guabiraba, Clare E. Bryant, Brian J. Ferguson

## Abstract

The sensing of viral nucleic acid by pattern recognition receptors (PRRs) is essential for initiation of a type-I interferon response against infection. Intracellular DNA sensing PRRs are responsible for initiating innate immune responses to poxviruses and other double-stranded DNA viruses. Poxviruses, in turn, encode an armoury of immunomodulators that inhibit this host defence mechanism. DNA-dependent protein kinase (DNA-PK) is an essential component of the cGAS/STING-dependent viral DNA sensing machinery that leads to the initiation of a type-I interferon response during poxvirus infection. Poxviruses counter this host sensing mechanism using the OPG020/OPG031 family of proteins that target DNA-PK, interfering with its ability to bind viral DNA. Although the DNA-PK complex, known also for its role in double strand break repair, is present across multiple taxa, its function in innate immunity outside mammals is unexplored. Here we analysed the contribution of DNA-PK to viral DNA sensing in chickens, a species that is evolutionarily distant from mammals, but that is also infected by poxviruses. We found that DNA-PK functions as a DNA sensor in chickens, and this process is countered by OPG020/OPG031 family members found in fowlpox virus. This host/pathogen interaction is conserved in a broader range of species than other mechanisms of poxvirus antagonism of innate immune sensing, which may reflect the difficulty of the host in evolving escape mechanisms that interfere with a protein that is essential for maintenance of genomic stability.

## Introduction

The molecular arms race between host and pathogen results in the evolution of increasing complexity in both host immune systems and pathogen effector mechanisms [1, 2]. This competition is exemplified by the interplay between host innate immune responses and antagonists of these systems deployed by viruses that provide extensive selective pressure, shaping the evolution of host immune responses [3, 4]. Across all vertebrate taxa, pattern recognition receptors (PRRs) have evolved to sense infection by directly binding molecular components of pathogens. PRR signalling activation results in the rapid transcription of antimicrobial genes, including interferons and cytokines in mammals. Viral effector mechanisms target PRR signalling to effectively block the production of these inflammatory mediators, allowing them to increase the temporal window for their replication and spread. Determination of the mechanisms of antiviral innate immunity in species that cross different orders of life has significant implications both for understanding the evolution of host/pathogen interactions and for understanding the outcome of zoonotic viral disease severity.

Intracellular PRRs that sense and respond to nucleic acids are of particular importance in initiating the host response to virus infection [5]. The DNA sensing PRRs initiate interferon transcription following DNA virus infections by sensing viral genomic DNA. A number of viral DNA sensors have been identified that are critical for defence against DNA viruses. These sensors, including cGAS and DNA-PK drive the STING-dependent IRF and NF-⍰B activation responsible for initiating an antiviral transcriptional programme [6]. Multiple families of DNA viruses are detected by PRRs that activate STING, including *poxviridae, herpesviridae* and *adenoviridae*, and these virus families encode multiple inhibitors of DNA sensing pathways that significantly reduce the production of interferons and cytokines from infected cells [7–9].

Poxviruses are an ancient virus family whose members can infect invertebrates, birds, reptiles and mammals. Poxviruses have rapid, lytic life cycles, necessitating extensive interactions with host innate immune systems. In mammals, poxviruses trigger cGAS and DNA-PK-dependent interferon production via STING [9–11] but encode an armoury of proteins that target this intracellular signalling pathway so effectively that the wild type virus elicits little to no interferon production from infected cells [9, 12–14]. The effective inhibition of viral DNA sensing PRRs is achieved by several families of proteins that each inhibit a different step in these signalling pathways. Vaccinia virus (VACV) is perhaps the best studied poxvirus in terms of immune evasion mechanisms [14–16]. VACV encodes several defined inhibitors of DNA PRRs including the OPG020/OPG031 family that binds the Ku70/80 component of DNA-PK and sterically impedes DNA-binding to reduce viral DNA-driven interferon production from infected cells [17–19]. The VACV OPG031 is also named C4L in the VACV Copenhagen strain nomenclature. The VACV OPG020 protein is C10L (Copenhagen) or C16L (Western Reserve strain nomenclature) where it encoded by two identical gene loci VACWR010 and VACWR209. VACV also encodes an enzyme, OPG188a (B2/poxin) that cleaves the second messenger cGAMP that is produced by cGAS and binds and activates STING [20, 21], and OPG067 (E5) that is reported to bind directly to cGAS and target it for degradation [22].

We previously defined the function of the chicken cGAS/STING signalling axis in the sensing of the avipoxvirus, fowlpox virus (FWPV) [12], confirming the conservation of this pathway in sensing mammalian and avian poxviruses. In this study we asked two specific questions. Firstly, given that bird genomes encode cGAS and DNA-PK, but not IFI16, we asked whether DNA-PK functions as an innate sensor of exogenous DNA and DNA viruses in birds as it does in mammals [10, 23, 24]. Secondly, we analysed whether this fundamental mechanism of intracellular viral sensing is targeted by poxviruses that infect birds. Our data illustrate the importance of DNA-PK in sensing FWPV and exogenous DNA in chicken cells and determines the breadth of conservation of the mechanism encoded by poxviruses to inhibit DNA-PK-dependent DNA sensing. We find that this mechanism of interfering with viral DNA sensing exists across more species than other known methods by which poxviruses block intracellular DNA PRRs, cementing DNA-PK as a key component of anti-poxvirus immunity across different orders of life.

## Material and Methods

### Reagents

Calf Thymus (CT) DNA (Sigma) and Herring Testes (HT) DNA (Sigma), polyinosinic-polycytidylic acid (poly(I:C), Invivogen), were diluted in nuclease-free water (Ambion, ThermoFisher).

### Plasmids

Chicken Ku70 and Ku80 were amplified from cDNA samples prepared from HD11 cells as described below. The amplified genes were cloned into pcDNA3.1 using the NEBuilder® HiFi DNA Assembly Cloning Kit (New England Biolab). A myc-tag and an HA-tag was added to the C-terminal of pcDNA3.1-Ku70 and Ku80, respectively, using the Q5® Site-Directed Mutagenesis Kit (New England Biolab). Codon optimised FPV020 and FPV006/255 open reading frames with a C-terminal FLAG-tag were ordered as gBlocks from IDT and cloned into pcDNA4. The sequences of all plasmids were verified by Plasmidsaurus Inc.

### Cell Culture

HD11 cells, an avian myelocytomatosis virus (MC29)-transformed chicken macrophage-like cell line, were incubated at 37°C, 5% CO2. They were grown in RPMI (Sigma-Aldrich, Germany) complemented with 2.5% volume per volume (v/v) heat-inactivated foetal bovine serum (FBS; Sera Laboratories International Ltd), 2.5% volume per volume (v/v) chicken serum (New Zealand origin, Gibco, Thermo Fisher Scientific), 10% Tryptose Phosphate Broth solution (Gibco, Thermo Fisher Scientific), 2 mM L-glutamine (Gibco, Thermo Fisher Scientific), 50 µg/ml of penicillin/streptomycin (P/S; Gibco, Thermo Fisher Scientific). Chicken embryonic fibroblasts (CEFs) (Pirbright Institute, Woking) were incubated at 37°C, 5% CO2 and were grown in Dulbecco’s Modified Eagle Medium (DMEM) -F12 with Glutamax (Gibco), 5% v/v FBS, and 50 µg/ml P/S. DF-1 cells were grown in DMEM with 10% v/v FBS, 2.5% v/v chicken serum and 50 µg/ml P/S. Chicken bone marrow derived macrophages (BMDM) were generated as previously described (17). Briefly, femurs and tibias of 4 week-old immunologically mature White Leghorn (PA12 line) outbred chickens were removed, both ends of the bones were cut and the bone marrow was flushed with RPMI supplemented with P/S. Cells were then washed and re-suspended in RPMI, loaded onto an equal volume of Histopaque-1077 (Sigma-Aldrich, Germany), and centrifuged at 400 g for 20 min. Cells at the interface were collected and washed twice in RPMI. Purified cells, pooled from three homozygous chickens, were seeded in triplicates at 1×10^6^ cells/ml in sterile 60 mm bacteriological petri dishes in RPMI supplemented with 10% FBS, 25 mM HEPES, 2 mM L-glutamine, P/S and 25 ng/ml recombinant chicken colony stimulating factor 1 (CSF-1) (Kingfisher Biotech, Inc) at 41°C and 5% CO2. Half of the medium was replaced with fresh medium containing CSF-1 at day 3. At day 6, adherent cells were harvested and cultured in RPMI supplemented with 10% FBS, 25 mM HEPES, 2 mM L-glutamine, and P/S prior to stimulation.

### HD11 knockout generation by CRISPR-Cas9

CRISPR guide RNAs were designed according to the *Gallus gallus PKRDC* (DNA-PKcs) and *XRCC6* (Ku70) sequences obtained from the Ensembl database (release 94), single guide (sg)RNA sequences were designed targeting exon boundaries. Genome editing of HD11 was performed using ribonucleoprotein (RNP) delivery. tracrRNA was mixed with the target specific sgRNA (Table 1), followed by an incubation at 95°C. To form the RNP complex, the tracrRNA/sgRNA mix was incubated with the Cas9 protein (IDT, Leuven, Belgium) and electroporation enhancer at 21°C. To generate knockout cells, 1×10^6^ cells per guide were electroporated with the corresponding RNP complex using Lonza Electroporation Kit V (Lonza). After 48 h, the cells were expanded for future experiments and their DNA were extracted using the PureLink Genomic DNA Kit (Thermo Scientific, Waltham, MA, USA). The knockout efficiency was evaluated by genotyping the polyclonal cell populations using MiSeq (Illumina) according to a published method. The successfully edited populations were diluted to a concentration of 0.5 cell/well and seeded in 96-well plates. Individual clones were sequenced by MiSeq and the confirmed knockout clones were expanded for experiments (Supplementary Figure 1).

**Table 1.**
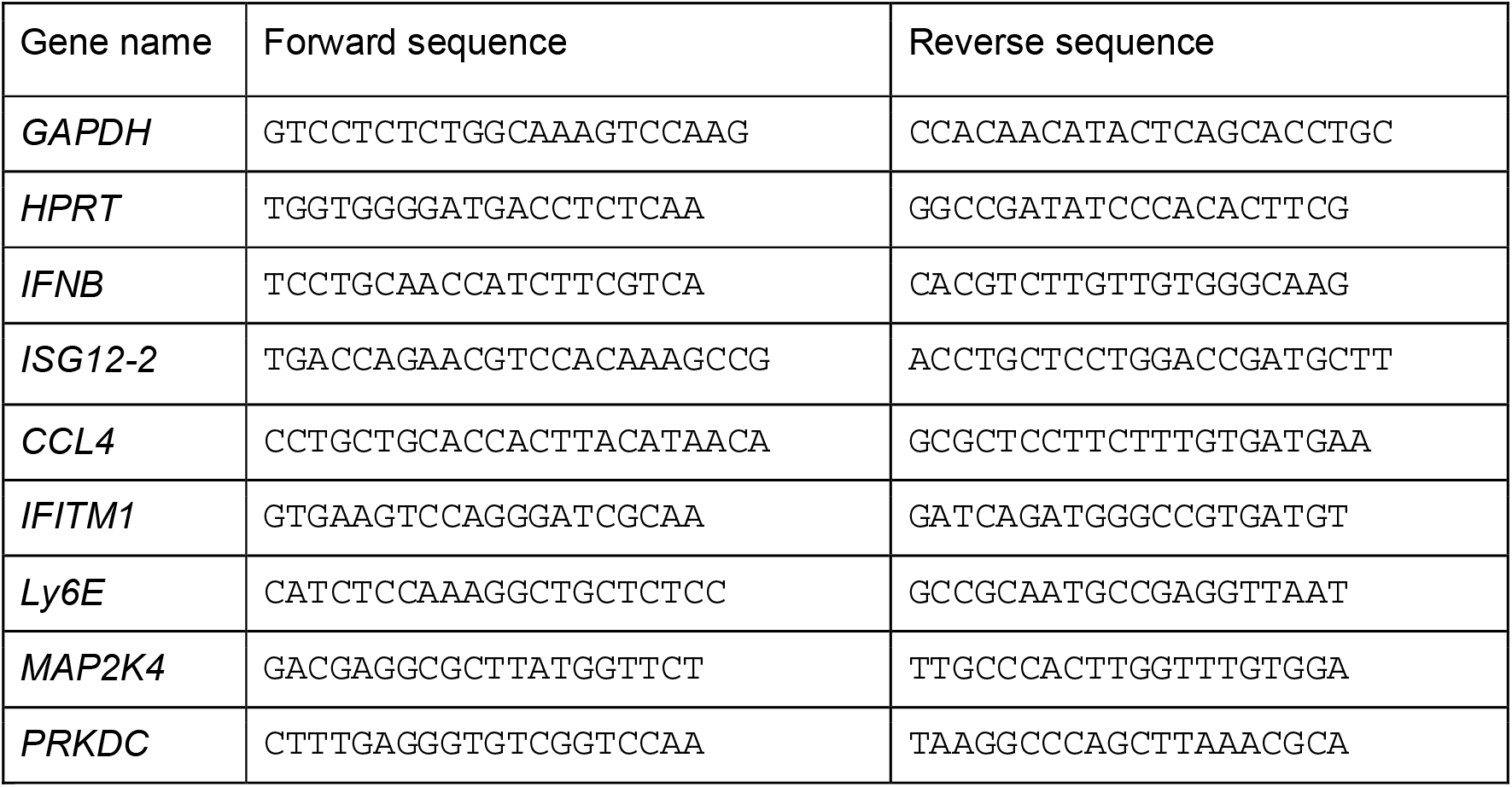
qRT-PCR primer sequences.

### Stimulation Assays

HD11 cells were seeded in 12-well plates at a density of 3×10^5^ cells/well. In the following day, the cells were transfected using TransIT-LT1 (Mirus Bio, USA) with CT-DNA (1, 2, or 5 µg/ml) or Poly(I:C) (1 µg/ml), and harvested 6 or 16 h post-transfection. In the priming assays, IFNα (200 ng/ml) was added 16 h prior to transfection.

### RNA Extraction

Cells were lysed by overlaying with 250 µl of lysis buffer containing 4 M guanidine thiocyanate, 25 mM Tris pH 7, and 143 mM 2-mercaptoethanol. As a second step, 250 µl of ethanol was added, and the solution was transferred to a silica column (Epoch Life Science, Inc., Sugar Land, TX, USA) and centrifuged; all centrifugation steps were performed for 90 s at 16,600 g. The bound RNA was washed by centrifugation with 500 µl of buffer containing 1 M guanidine thiocyanate, 25 mM Tris pH 7, and 10% ethanol, followed by a double washing step with 500 µl of wash buffer 2 [25 mM Tris pH 7 and 70% (v/v) ethanol]. RNA was eluted by centrifugation in 30 µl of nuclease-free water and the concentration was measured using a NanoDrop 2000 Spectrophotometer (Thermo Scientific, Waltham, MA, USA).

### cDNA and qPCR

Using 500 ng of RNA extracted from HD11 cells, cDNA was produced using SuperScript III reverse transcriptase, following the manufacturer’s protocol (Thermo Scientific, Waltham, MA, USA). Samples were diluted in nuclease-free water in a 1:2.5 ratio. One μl of the diluted product was used for quantitative PCR (qPCR) in a final volume of 10 μl. qPCR was performed using SybrGreen Hi-Rox (PCR Biosystems Inc.) using primers described in Table 1. Fold change in mRNA expression was calculated by relative quantification using hypoxanthine phosphoribosyltransferase (HPRT) as endogenous control.

### Fowlpox Virus Growth and Titration

FWPV WT (FP9) and mutants were propagated in primary chicken embryonic fibroblasts (CEFs) and grown in DMEM-F12 (Thermo Fisher Scientific, Waltham, MA, USA) containing 1% FBS and 5% P/S, and harvested 5 days later. Ten-fold dilutions of cell supernatants were prepared in serum-free DMEM-F12 and used to inoculate confluent monolayers of CEFs for 1.5 h at 37°C. Cells were then overlaid with 2xMEM: CMC (1/1 ratio). The plaques were counted seven days later after fixation. For infection experiments HD11 cells were seeded in 12-well plates in the day prior to infection. Fowlpox viruses were diluted in serum-free DMEM-F12 at a multiplicity of infection (MOI) of 3 and added in the cells (1 ml per well). Infected cells and supernatants were collected from infections at 16h and 24h post-infection.

### Homology searching and databases

Identification of PRR pathway inhibitors in poxviridae. A broadly representative panel of 54 complete reference poxvirus genomes was selected from the Bacterial and Viral Bioinformatic Resource Center (https://www.bv-brc.org). Vaccinia sequences for the protein families of interest were searched against predicted proteomes derived from this genome panel using BLASTp at low stringency (e-value cut off 0.1, max hits 500). Putative homologues were confirmed by reciprocal best BLASTp into the vaccinia proteome, followed by sequence alignment and, where relevant, examination of predicted domain structure. A further, iterative, HMM-based search strategy was employed to ensure that no detectable homologues had been missed. Using the vaccinia sequences as the seed, iterative HMMER (http://hmmer.org) searches were carried out against the entire uniprotKB database. These searches were run until convergence or divergence and results filtered to identify poxviridae derived sequences. These searches returned no additional homologues from the genome panel initially selected suggesting that our initial search had captured all identifiable homologues.

### Identification of DNA-PK subunits in chordates

A representative panel of chordate genomes with a focus on groups widely associated with poxvirus infection (reptiles, birds, mammals) was selected (Supplementary Table 1). Human DNA-PKcs, Ku70, and Ku80 were searched against predicted proteomes from these genomes using BLASTp with default settings. Putative homologues were validated via reciprocal BLASTp into the human proteome. Where no homologous proteins were identified within a taxa a phylogenetically close taxa was also searched in order to reduce the possibility of false negatives. In all cases where we failed to identify protein homologues within a genome the same was true of sister taxa.

### Phylogenetic reconstruction of OPG020/OPG031 family

Phylogenetic analysis was carried out as previously described [25]. Briefly, Sequences were aligned in Jalview with MAFFT (L-INS-I) [26, 27]. Informative sites identification and alignment trimming was carried out with TrimAI (Strict settings) [28]. Subsequent substitution model selection (ModelFinder[29]) and phylogenetic reconstruction was carried out in IQ-Tree.

### Co-Immunoprecipitation

2 million DF-1 cells were seeded in 60mm dish one day prior to transfection. Each dish was transfected with 1.5 µg each of chicken Ku70 and Ku80, along with 1 µg of GFP, FPV020 or FPV006/255 using FuGENE® HD (Promega) at a ratio of 1 µg plasmid to 3 µl of reagent. After 24 hours, transfected cells were washed once with ice cold PBS before being lysed in RIPA buffer (50mM Tris-HCl pH8.0, 150mM NaCl, 1% NP-40, 0.1% SDS, 0.5% Sodium deoxycholate) supplemented with cOmplete™ Protease Inhibitor Cocktail (Roche). Lysate were spun for 10 minutes at 20,000 g and the clear lysate were harvested. 10% of samples were kept as input and the rest were subjected to incubation with 20 µl of Anti-FLAG® M2 Magnetic Beads (Millipore) overnight with rotation at 4°C. The beads were washed 3 times with TBS before being eluded in SDS-PAGE sample buffer.

### Immunoblotting

Samples were boiled at 98°C before being separated by a 10% Tris-glycine gel. Proteins were transferred on 0.2-μm nitrocellulose membranes (Cytiva) using the Trans-Blot Turbo Transfer System (Bio-Rad). Membranes were blocked with 5% BSA in TBS-T for 1 hour before being incubated with the following primary antibodies: anti-myc-tag at 1:2000 (16286-1-AP, Proteintech), anti-HA-tag at 1:2000 (81290-1-RR, Proteintech), anti-Flag-tag at 1:2000 (F7425, Sigma) and anti-Actin at 1:2000 (ab8226, Abcam). The membranes were incubated with secondary antibody anti-Rabbit IgG 680nm at 1:10,000 (926-68071, Licor) before being imaged on an Odyssey CLx Imager (Licor).

### Microarray/RNAseq

The FWPVΔ020 deletion virus was previously described [30, 31]. To investigate host responses to these mutants, primary chicken embryo fibroblasts (CEF) were infected with FWPVΔ020 incubated for 16 hours post-infection. Total RNA was extracted, and microarray-based transcriptional profiling was performed using Affymetrix GeneChip Chicken Genome Arrays, which contain probes for both FWPV and host genes. The microarray data were deposited in the ArrayExpress database (E-MTAB-7276), along with transcriptomic datasets from other FWPV deletion mutants. Raw data files in .CEL format are publicly available, with systematic sample processing conducted to mitigate batch effects. The original microarray data used in this study have been previously published [13] and deposited in accordance with MIAME guidelines in the public database ArrayExpress (http://www.ebi.ac.uk/microarray-as/ae/; Acc. No: E-MTAB-7276). Data were reanalyzed for this study at both the probe and gene levels using the Transcriptome Analysis Console (TAC) 4.0 software from Affymetrix. Microarray data were analysed with default settings, where a gene was considered expressed in a particular condition if it was detected in 50% or more of the samples with a Detection Above Background (DABG) *p*-value of less than 0.05. Additionally, a threshold of a 1.5-fold change in expression was applied to identify differentially expressed genes.

### Pathway analysis

Differentially regulated genes underwent Gene Ontology (GO) analysis, which classified associated mechanisms into biological processes (BP) and dot plot visualisation was done using ShinyGO (v0.80; [32]). A False Discovery Rate (FDR) *p*-value threshold of <0.05 was used as a standard metric to quantify the most closely related GO terms. The heatmap illustrating gene expression patterns was generated using the seaborn library (v0.13) for advanced visualisation in Python 3.8+, with additional customization provided by matplotlib (v3.9.0).

### Protein structure prediction

AlphaFold2 (AF) was used for predicting the structures of FWPV006 and FWPV020 and AF2-Multimer for Gallus Gallus Ku70/80 heterodimer [33]. The top ranked hit for each was assessed for its confidence score and coloured according to AF prediction confidence. The dimeric assemblies of FWPV006 and FWPV020 were created by docking the N- and C-terminal domains onto the structure of C16 (C10) due to their similarity and the inability of AF to correctly predict these accurately. All structural figures including electrostatics were made using ChimeraX [34].

### Statistical analysis

Prism 9 (GraphPad) was used to generate graphs and perform statistical analysis. Data were analysed using an unpaired t test with Welch’s correction unless stated otherwise. Data with P < 0.05 was considered significant and 2-tailed P-value were calculated and presented as: *p < 0.05, **p < 0.01, ***p < 0.001; ****p < 0.0001. Each experiment has at least two biological replicates unless stated.

## Results

### DNA-PK positively regulates innate immune viral DNA sensing in chickens

DNA-PK consists of three subunits, DNA-PK catalytic subunit (DNA-PKcs, PRKDC/XRCC7), Ku70 (XRCC6) and Ku80 (XRCC5) functioning together as a heterotrimeric protein complex. Broad evolutionary analyses have identified homologues of DNA-PKcs, Ku70, and Ku80 across the eukaryotes and limited functional analysis in diverse lineages supports a conserved role in non-homologous end joining (NHEJ) double-strand DNA break repair [35–37]. These genes are therefore important for the maintenance of genomic stability and, in jawed vertebrates, for V(D)J recombination [38]. By sensing viral DNA, DNA-PK has another established function in innate immunity in humans and mice as a positive regulator of virus-driven IFNβ production [10, 24, 39], although other reports describe a function of the kinase activity of DNA-PKcs in negatively regulating this process [40]. In part to address this disparity, and as the degree of conservation of this DNA-PK’s function in antiviral responses across taxa is unexplored, we first examined the distribution of the DNA-PK complex within the chordates. This order contain the natural hosts of the economically important and human infective chordopoxvirus family. We sampled a range of chordate genomes covering this lineage, with a focus on major chordopoxvirus targets (birds, reptiles, mammals). The DNA-PK complex was identified in the majority of genomes examined and, in every bird, reptile, and mammal that we searched (Figure 1A). If the innate immune function of the DNA-PK complex is also conserved, as shown for humans and mice, then targeting this activity may also be important for poxvirus lineages beyond vaccinia virus. With this in mind, we next examined the specific target of the vaccinia virus OPG020 protein, the Ku80 subunit of the DNA-PK complex. Ku80 shows high sequence level conservation across the chordates, particularly in the DNA binding domain, likely reflecting its important function in DNA repair (Figure 1B). We also note that this DNA binding domain is the target of OPG020 binding.

**Figure 1.**
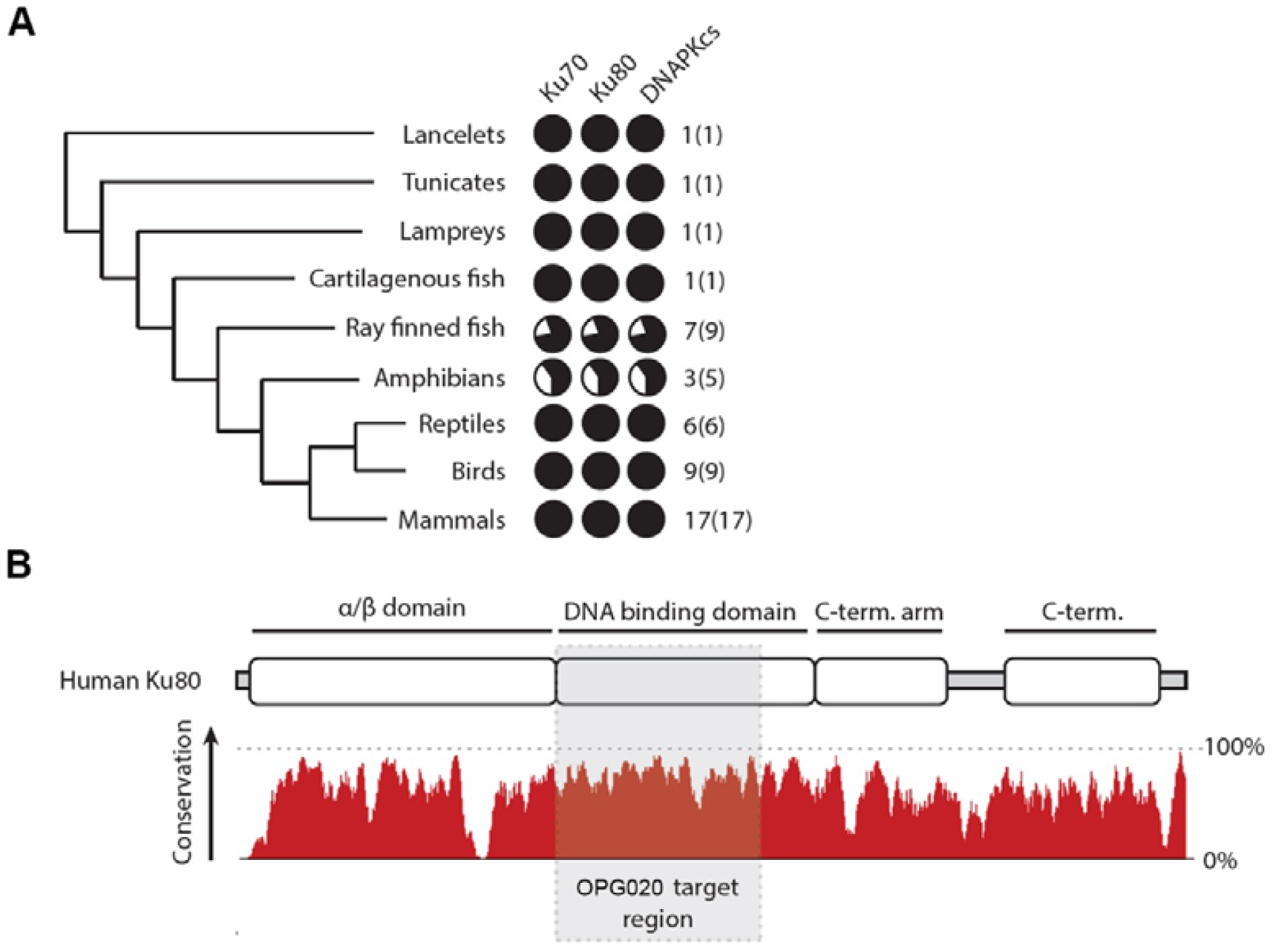
Conservation of the DNA-PK complex. A) Degree of conservation of Ku70, Ku80 and DNA-PKcs genes across chordates. Numbers indicate the presence of genes found in each taxonomic group compared with the number of species searched. B) Amino-acid level conservation of Ku80 across chordates. Domain structure of Ku80 is shown above and amino-acid percentage conservation shown below. The region of Ku80 targeted by VACV OPG020 is shown in grey.

The Ku proteins and DNA-PKcs carry out NHEJ functions in chicken cells [41] but they have not been studied in the context of avian innate immune responses. To functionally assess the contribution of DNA-PK to intracellular DNA sensing in chickens we used the HD11 chicken macrophage cell line that has an active STING-dependent signalling pathway [12] to generate DNA-PKcs knockout lines using CRISPR/Cas9 and confirmed the generated indels by sequencing across the target loci (Supplementary Figure 1A, B). Attempts to produce Ku70 knockout were unsuccessful as, despite multiple sgRNA guides targeting different exons being used, very few stable clones could be recovered, and those that were recovered were found to have high background levels of cytokine and chemokine transcription that rendered the assessment of Ku70’s role in DNA sensing intractable in this system.

Concentrating on DNA-PKcs, we found this gene (*PRKDC*) to be transcribed in both chicken bone marrow derived macrophages and HD11 cells, and that its transcription is upregulated by interferon alpha (IFNα) treatment (Figure 2A). We were able to generate multiple confirmed DNA-PKcs knockout clonal cell lines in HD11 that, in the resting state, did not have altered inflammatory gene transcription compared to parental lines. In these cells the DNA-dependent transcription of type-I interferon (*IFNB*) and *ISG12*.*2* was significantly reduced when compared to WT controls (Figure 2B, C). The transcription of CCL4, however, was unaffected by loss of DNA-PKcs (Figure 2D). In comparison, no defects were found in RNA-driven transcriptional responses in DNA-PKcs knockout cells (Figure 2E) consistent with a DNA-specific role for DNA-PK in intracellular nucleic acid sensing in chickens.

**Figure 2.**
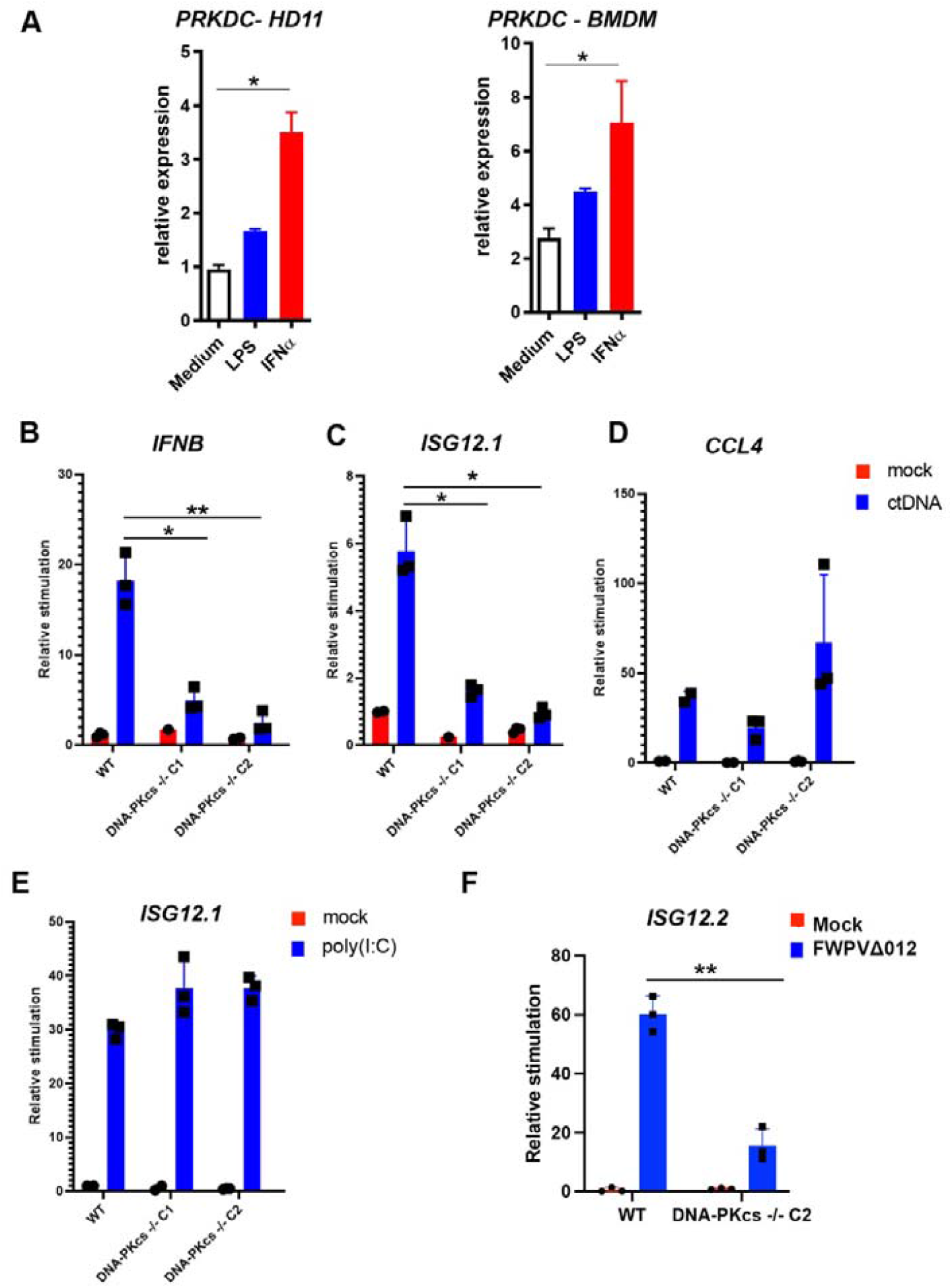
Exogenous DNA and Fowlpox virus sensing are impaired in DNA-PKcs knockout chicken cells. **A)** HD11 or BMDM cells were treated with 10 nM LPS or 50 µM IFN⍰ for 6 hours and the level of *PRKDC* mRNA (encoding the DNA-PKcs protein) was measured by RT-qPCR. p<0.01, n=3. Data is presented as mean ± SEM and analysed by two-tailed Student’s T test with n = 3; ⍰p < 0.05. WT or DNA-PKcs^-/-^ HD11 cells were stimulated with CT-DNA (5 µg/ml) and transcription of B) *IFNB*, C) *ISG12*.*2* and D) *CCL4* were measured by qRT-PCR 6 h later. E) WT or DNA-PKcs^-/-^ HD11 cells were stimulated with Poly(I:C) (2 µg/ml) and and transcription of *ISG12*.*2* was measured by qRT-PCR 6 h later. F) WT or DNA-PKcs^-/-^ cells were infected with FWPVΔ012 and *ISG12*.*2* was measured by qRT-PCR 24 h later. Data is presented as mean ± SEM and analysed by two-tailed Student’s T test with n = 3; ⍰p < 0.05, ⍰⍰p < 0.01.

Wild-type poxviruses, including FWPV, often fail to induce an innate immune response due to the presence of a large number of intracellular PRR inhibitors [9, 30]. Using a FWPV strain FP9 lacking an immunomodulator (FPV012) that inhibits IFN-I production [13], we previously demonstrated that cGAS and STING are essential for sensing FWPV infection and that transcription of *ISG12*.*2* is a cGAS/STING-dependent signalling output in chicken cells [12]. Here we infected WT and DNA-PKcs^-/-^ macrophage cell lines with FWPVΔ012 strain and observed lower induction of *ISG12*.*2* transcription in DNA-PKcs^-/-^ cells compared with WT cells (Figure 2F). This data indicates that DNA-PKcs acts in concert with cGAS and STING in sensing FWPV infection in chicken cells in the same way that it does in mammals to sense VACV [10].

Chicken cells use a STING/TBK1/IRF7 pathway to sense DNA and FWPV infection To further define the DNA-PK-dependent innate sensing pathway in chicken macrophages we used IRF7 knockout cells HD11 cells [25] and generated clonal TBK1 HD11 knockout lines (Supplementary Figure 1). In mammals, following DNA stimulation, TBK1 phosphorylates STING and IRF3 allowing dimerisation nuclear translocation of IRF3 and subsequent IFN-I transcription. Chickens lack IRF3 but express an IRF7 with a C-terminus similar to mammalian IRF3 that has a distribution of serine residues that mimic the mammalian TBK1 phosphorylation sites [25]. Knockout of TBK1 or IRF7 in HD11 cells resulted in a loss of DNA-driven *IFNB* and *ISG12*.*2* transcription (Figure 3A). Similarly the transcription *ISG12*.*2* in response to FWPVΔ12 infection was completely dependent on TBK1 and IRF7 (Figure 3B). These data, combined with our prior description of cGAS and STING knockout macrophages [12] indicate that chickens sense cytoplasmic DNA and FWPV infection via DNA-PK, and that this uses a canonical cGAS/STING/TBK1 pathway to drive IFN-I transcription with IRF7 as the dominant transcription factor. As such these core components of the intracellular DNA sensing pathway are functionally consistent between mammals and birds.

**Figure 3:**
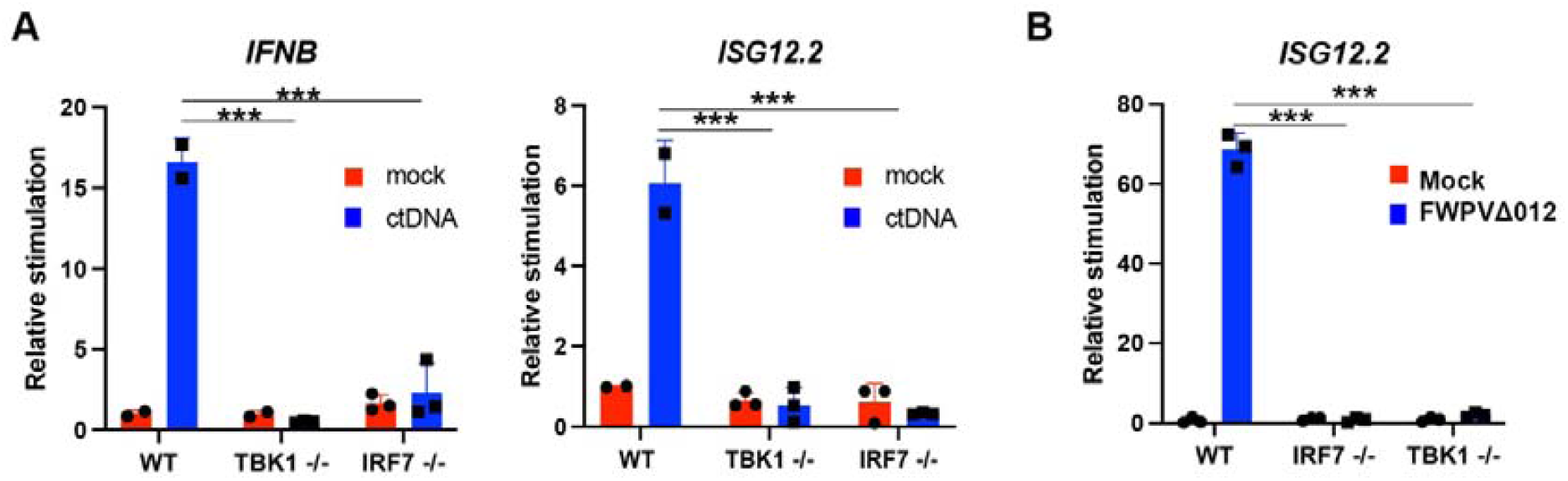
Fowlpox sensing by DNA-PKcs / TBK1 / IRF7. A) WT, TBK1^-/-^ or IRF7^-/-^ HD11 cells were stimulated with CT-DNA (5 µg/ml) and transcription of *IFNB* and *ISG12*.*2* were measured by qRT-PCR 6 h later. B) WT, TBK1^-/-^ or IRF7^-/-^ HD11 cells were mock infected or infected with FWPVΔ012 and *ISG12*.*2* was measured by qRT-PCR 24 h later. Data is presented as mean ± SEM and analysed by two-tailed Student’s T test with n = 3; ⍰⍰***p < 0.01.

The poxvirus OPG020/OPG031 family are present across multiple poxvirus genera Poxviruses use a plethora of immune modulators to inhibit PRR signalling and inhibit interferon production [14]. The OPG020 and OPG031 proteins made by vaccinia virus bind directly to Ku and inhibit its ability to sense viral DNA [17–19]. Since the innate sensing function of DNA-PK is conserved across mammals and birds we postulated that the immune evasion mechanism used by the OPG020/OPG031 poxvirus family might also be. We therefore used protein sequence homology searches to identify OPG020/OPG031 homologues and analyse their distribution across a broadly representative selection of poxviruses (Figure 4A and [42]). As previously reported, the B2/poxin family demonstrated the broadest distribution, being identified in mammalian and insect infective (entomopoxvirus) poxviruses [43]. However, the distribution of B2/poxin was extremely sparse beyond the orthopoxviruses and centapoxviruses. Of the other inhibitors examined, the OPG020/OPG031 family appeared the oldest, being present in the relatively early branching avipoxvirus family. Further examination of the relative positions of OPG020/OPG031 family genes within their respective genomes revealed that this family is almost exclusively localised to the genomic termini, either within or adjacent to the inverted terminal repeat (ITR) regions. These genomic regions are highly enriched in virulence genes, suggesting that a virulence mediating function may be conserved across the OPG020/OPG031 family (Figure 4B). This analysis of the conservation of OPG020/OPG031 family homologues indicates that this family is present across more poxvirus genera than the OPG188a and OPG067 families (Figure 4A) and suggest therefore that targeting DNA-PK may represent a broadly conserved and ancient strategy for poxvirus immune evasion, likely reflected in the consistent positioning of OPG020/OPG031 family genes within ITRs and genome termini.

**Figure 4.**
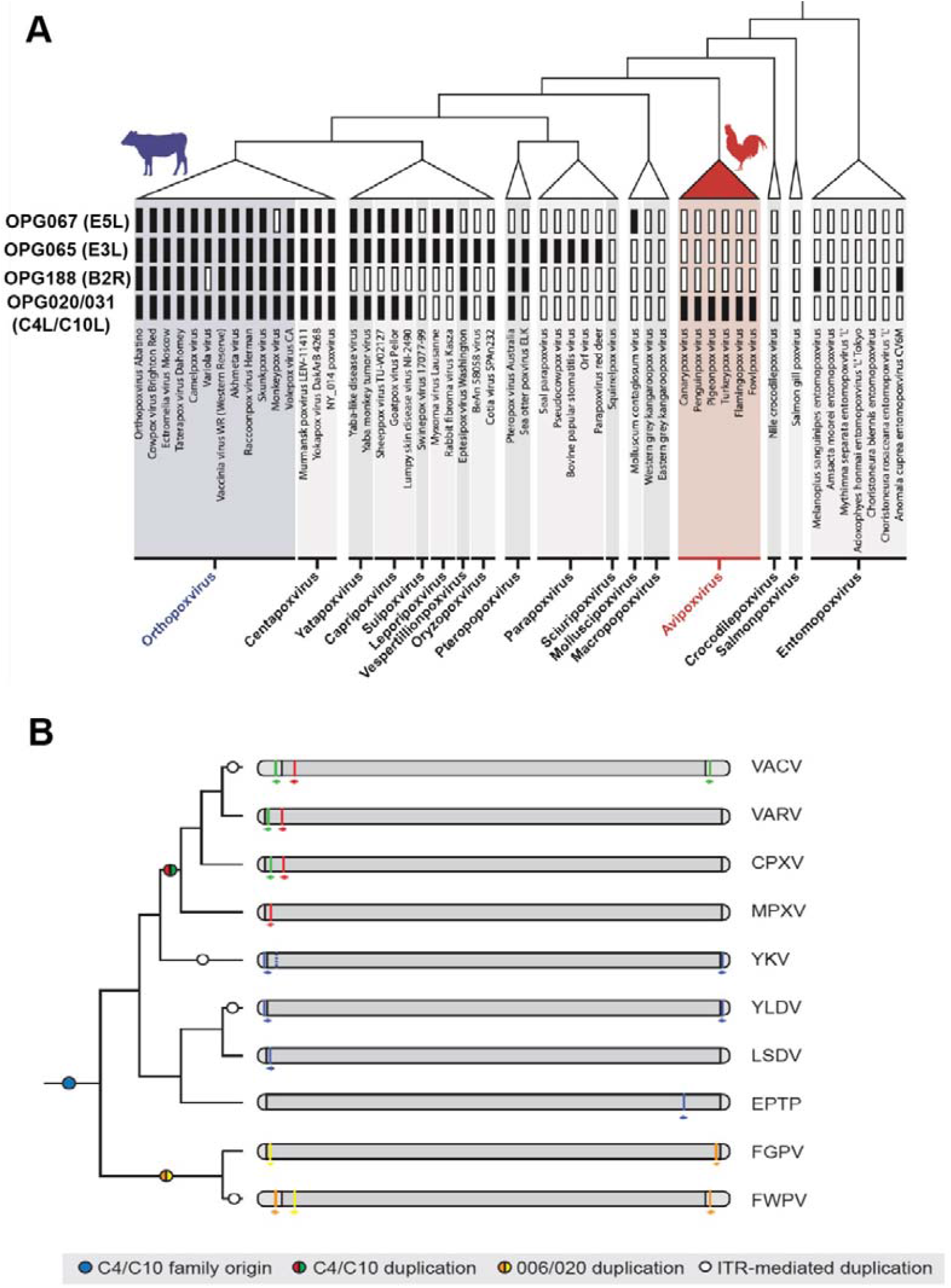
The OPG020/OPG031 family of poxvirus immunomodulators is present in multiple poxvirus genera. A) Gene-level conservation of poxviral immunomodulators OPG020, OPG031, OPG188a, OPG065 and OPG067 and across poxviridae. B) Genomic locations of OPG020/OPG031 family genes in selected poxviruses. Coloured bands represent individual gene positions, arrows denote direction of open reading frame.

We identified three OPG020/OPG031 family member genes in FWPV. This family contains a characteristic bi-domain structure (Figure 5A). The N-terminal domain is a structural mimic of the prolyl-hydroxylase family responsible for regulation of hypoxia-inducible factor 1⍰ (HIF-1⍰) [44], while the C-terminal domain binds Ku70/80, blocking DNA binding to DNA-PK [18]. The C-terminal domain amino-acid sequence contains a distinctive ‘LCYC’ motif that is very highly conserved across family members and is structurally essential for the domain fold [17, 19]. Two of these genes, 006 and 255, encode identical proteins and, like VACV OPG020, are duplicate genes found in the inverted terminal repeats (ITR) of vaccinia that contain multiple duplicated genes. The third gene in FWPV is 020 which has a divergent sequence from 006/255 but is still a member of the same family. To assess further the relationship between the poxviral OPG020/OPG031 family members we carried out a phylogenetic analysis of family members (Figure 5B). This analysis indicates that there have been multiple independent gene duplication events during the evolution of poxviruses that have led to many poxviruses carrying more than one copy of these genes. The duplication events, for example, that led to the appearance of OPG031 and OPG020 in VACV occurred independently of the event that led to 020 and 006/255 in FWPV. This can be further evidenced by the presence of a C-terminal extension in all three FWPV proteins (006, 020, 255) that is not present in any of the orthopoxvirus family members (Supplementary Figure 2). As such there is a clear evolutionary pressure on most poxvirus families to maintain at least one copy, and in many cases multiple copies, of the OPG020/OPG031 family.

**Figure 5.**
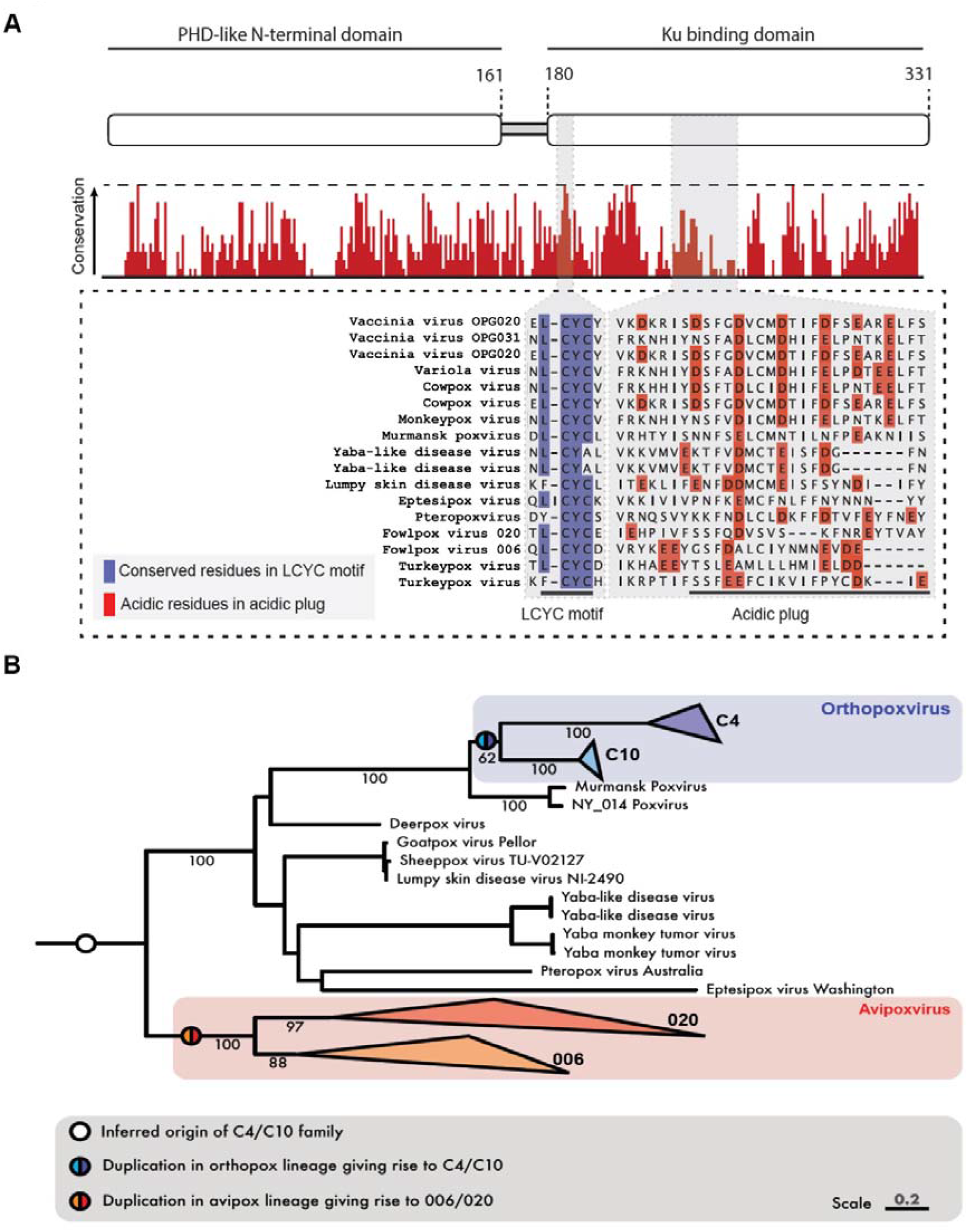
Conservation of the OPG020/OPG031 family. A) Amino-acid level conservation across the OPG020 protein. Domain structure of OPG020/OPG031 (top) with % amino acid conservation below, specifically highlighting the highly conserved LCYC motif and the variability across the acidic plug that binds Ku. B) Phylogenetic tree of OPG020/OPG031 family, branch labels represent ultrafast bootstrap support at key nodes.

Whilst it is true that the OPG020/OPG031 family is well maintained in terms of presence, examination of sequence level conservation across family members demonstrated a notable lack of strict sequence level conservation. The previously identified, highly conserved LCYC motif was indeed present across all identified genes, however the region of VACV OPG020 responsible for interacting with Ku80 and thus conferring inhibitory activity appeared only modestly conserved (Figure 5A). This prompted us to investigate experimentally the activity of the divergent avipoxvirus OPG020/OPG031 family proteins against DNA-PK from their natural host.

FPV020 but not FPV006/255 binds to chicken Ku70/Ku80 The structure of VACV OPG020 bound to human Ku has been solved by CryoEM [17]. In this structure, a OPG020 homodimer was shown to engage the Ku70/80 heterodimer DNA binding region, specifically at the electropositive surface created by the dimer interface between Ku70 and Ku80 (Figure 6A, PDB:8AG5). Several loops in the C-terminal domain of OPG020 dimer come together to form a structure that binds the inner surface of the Ku DNA-binding ring (Figure 6A). Primary sequence analysis of the OPG020/OPG031 family, including the FWPV proteins 020 and 006, indicates that the amino-acids forming the Ku-binding loop are not conserved across virus species (Figure 5A), suggesting the possibility of a different Ku-binding capacities across different poxviruses. To assess this further, we first used AlphaFold (AF) to predict three-dimensional structures of the chicken Ku 70/80, FPV020 and FPV006/255 proteins (Figure 6B). Both the N-terminal and C-terminal domains were predicted with high accuracy, but lower confidence was predicted in the loop between these domains and at the far C-terminus. The predicted structures of FPV020 and FPV006/255 suggest a consistent structure between these proteins and OPG020/OPG031. Dimeric assemblies of 020 and 006 were created by docking onto the VACV OPG020 structure, which overlays with very high confidence, indicating the potential for a conserved overall fold of these proteins.

**Figure 6.**
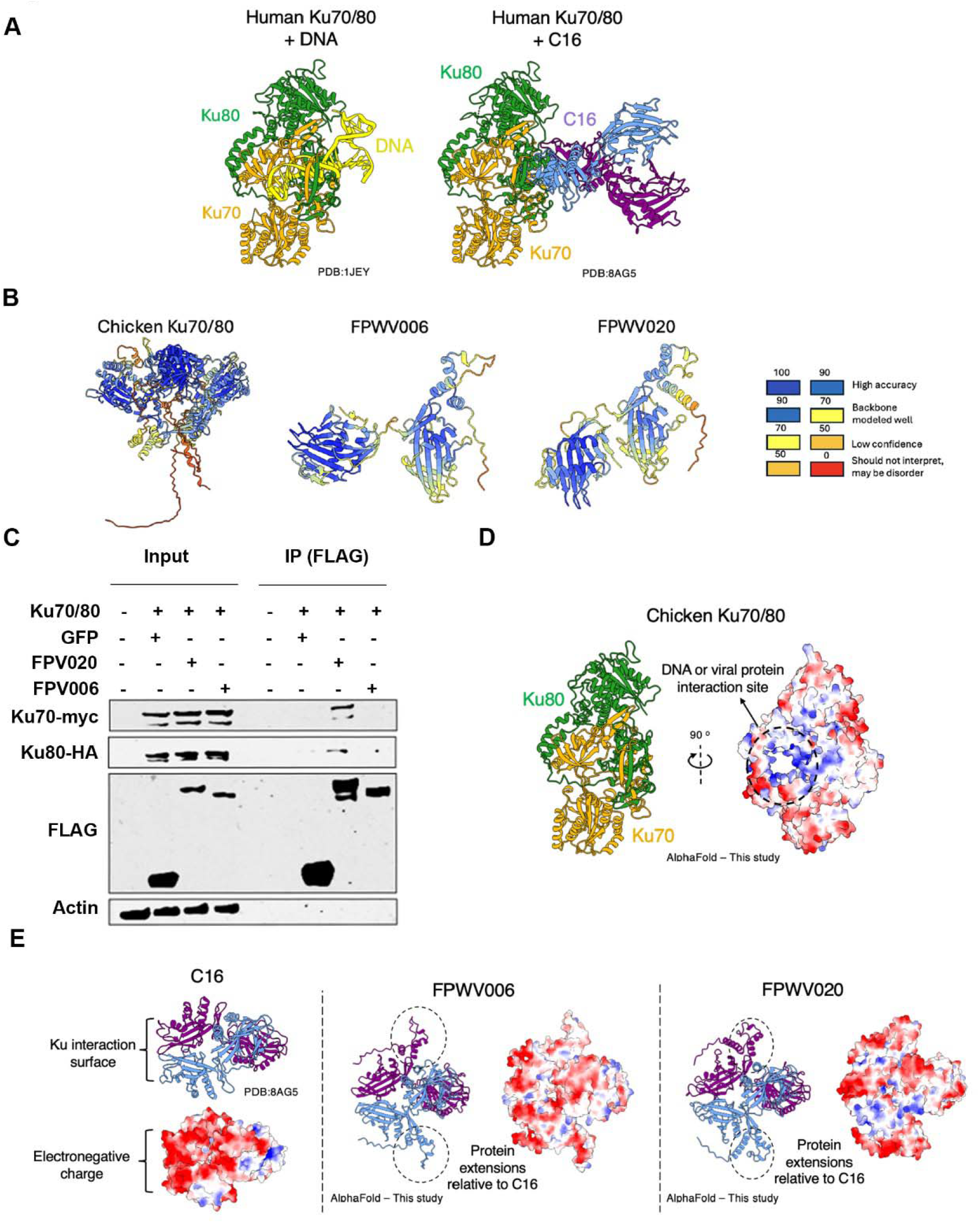
FPV020 binds chicken Ku70/80. A) X-ray crystallography structure of human Ku70/80 with DNA (PDB:1JEY) (Walker, 2001), Ku70 in orange, Ku80 in green and DNA in yellow (left). Cryo-EM structure of human Ku70/80 with VACV WR C16 (C10) (PDB:8AG5) (Rivera-Calzada et al., 2022), Ku70 in orange, Ku80 in green, C16 (C10) in blue and purple (right). B) AlphaFold-Multimer prediction of chicken Ku70/80, FPV006 and FWPV020 coloured according prediction confidence (right). C) Co-immunoprecipitation of FPV020 with chicken Ku70/80 in DF1 cells. Cells were transfected with indicated plasmids, immunoprecipitated with anti-FLAG beads and immunoblotted with the indicated antibodies. FPV020, but not FPV006 was co-immunoprecipitated with chicken Ku70/80. D) AlphaFold prediction of Chicken Ku70/80 (orange/green) and an electrostatic surface representation showing the electropositive patch in the DNA-binding region. E) Structures of C16 (C10) (PDB:8AG5), AlphaFold models of FWPV006 and FWPV020 in purple and blue with electrostatic surface comparisons.

To test the predicted interactions experimentally we used co-immunoprecipitation experiments in chicken DF1 cells. Due to the lack of cross-reactive antibodies against chicken Ku70/Ku80 proteins we initially cloned these open reading frames (ORF) as myc- and HA-tagged constructs and co-expressed them along with individual FLAG-tagged viral ORFs (FPV020 and FPV006/225). In these experiments FPV020 could pull down chicken Ku70/80 (Figure 6C), indicating a conserved function and mechanism of action between FPV020 and VACV OPG020. FPV006/255, however, did not co-precipitate with chicken Ku70/80 indicating a lack of conservation of interactions between the FPV020 and FPV006/255.

A likely explanation for this difference in binding lies in variations between FPV020 and FPV006/255 within the loop regions that extends into the central DNA-binding channel of Ku70/80. Chicken Ku70/80 is predicted to mimic the mammalian DNA-binding modality, having an electropositive ring that binds the electronegative DNA backbone (Figure 6D). This electropositive patch is utilised primarily for DNA binding, however the OPG020/OPG031 protein family have exploited this by creating proteins with high electronegativity to bind here and prevent DNA binding. VACV OPG020 has a strongly electronegative patch in this Ku-binding region (Figure 6E) and this is mimicked in the loop regions of FWPV020 that are predicted to interact with chicken Ku70/80 (Figure 6E). FPV020 contains a substantially long loop spanning residues 333–343 (FPV020 numbering). This extended loop is enriched in acidic residues, including Asp334, Glu335 and Glu338, which generate a strongly negatively charged surface. The conservation of this charge-distribution provides an explanation for the conservation of function of this family of proteins in the absence of strict primary sequence similarity conservation in the functional loop regions.

FPV006/255, however, possesses considerably shorter loop regions, particularly corresponding to residues 310–315 (OPG020 numbering). This hypothesis is further supported by electrostatic surface representations, which reveal differences in charge distribution between the two proteins. FPV020 presents a more extensive negatively charged surface within the putative DNA-binding interface than FPV006/255. Together, these observations provide a plausible structural explanation for why FPV020 interacts with Ku70/80, whereas FPV006/255 does not (Figure 6E).

### Fowlpox 020 is an immunomodulatory gene

To assess the conservation of function in the fowlpox C4/C10 homologues, we used an FWPVΔ020 deletion virus and analysed previously generated microarray data from primary chicken embryonic fibroblasts (CEF) infected with these viruses [13, 30]. We compared gene expression profiles of mutant strains to the wild-type fowlpox virus (FP9 strain), known for its disruption of DNA signalling pathways and suppression of interferon signalling in host cells [12, 30, 45]. Gene Ontology (GO) biological function analysis revealed significant differences in both knockout viruses compared to the FP9 strain, particularly in cell-to-cell signalling pathways (Figure 7A). We then screened the gene lists generated from the comparison between the knockout viruses and the wild-type virus for interferon and DNA-sensing immune response-related and cellular signalling transcripts [12, 46]. This screening revealed that several genes, including IFITM1, LY6E, MB21D1, IL6, FIB, TLR2-1, IFI30, and TNFAIP2, were subtly but significantly upregulated in one or both of the knockout mutants compared to the wild-type FP9 FWPV strain (Figure 7B).

**Figure 7.**
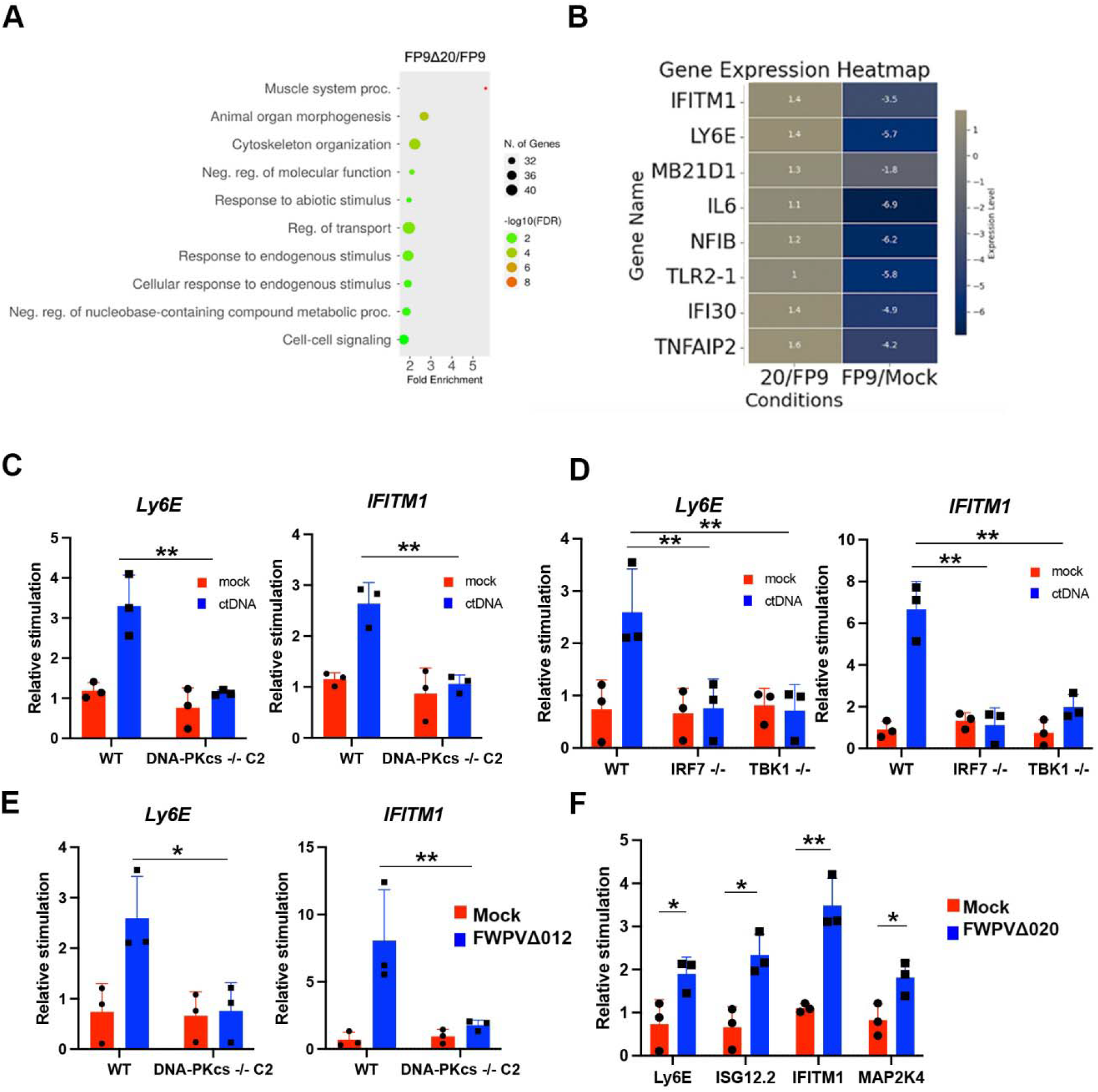
FPV 020 negatively regulates the IFN-I response during FWPV infection. A and B) Gene Ontology (GO) analysis using ShinyGO identified the top 10 enriched pathways in comparisons between FP9Δ06 vs. FP9 and FP9Δ020 vs. FP9, depicted as dot plots (Figures A and B, respectively). Dot size reflects the number of genes, and colour indicates the significance level (-log10(FDR)). C) A heatmap illustrating selected immune gene expression levels across conditions FWPVΔ006 vs. FP9, FWPVΔ020 vs. FP9, and FP9/Mock, generated using Python’s pandas, seaborn (v0.13), and matplotlib (v3.9.0). Axes display comparisons and gene names, while a colour bar provides a scale for gene expression levels. D) WT or DNA-PKcs^-/-^ HD11 cells were stimulated with CT-DNA (5 µg/ml) and transcription of *Ly6E* and *IFITM1* were measured by qRT-PCR 6 h later. E) WT or DNA-PKcs^-/-^ cells were mock infected or infected with FWPVΔ012 and *Ly6E* and *IFITM1* were measured by qRT-PCR 24 h later. F) WT, TBK1^-/-^ or IRF7^-/-^ HD11 cells were stimulated with CT-DNA (5 µg/ml) and transcription of *Ly6E* and *IFITM1* were measured by qRT-PCR 6 h later. G) HD11 cells were mock infected or infected with FWPVΔ020 and *Ly6E, ISG12*.*2, IFITM1* and *MAP2K4* were measured by qRT-PCR 24 h later. Data is presented as mean ± SEM and analysed by two-tailed Student’s T test with n = 3; ⍰p < 0.05, ⍰⍰p < 0.01.

The identification of interferon-stimulated genes *IFITM1* and *Ly6E* as 020-regulated genes in FWPV-infected CEFs is consistent with the function of these genes as immunomodulatory proteins and regulators of FWPV-driven interferon production. To identify the upstream signals responsible for transcription of *IFITM1* and *Ly6E* we analysed DNA-stimulated WT and DNA-PKcs^-/-^, TBK1^-/-^ and IRF7^-/-^ HD11 chicken macrophage cell lines. The DNA-driven induction of *IFITM1* and *Ly6E* transcription was dependent on DNA-PKcs, TBK1 and IRF7 (Figure 7C, D). Similarly, infection of these cells with FWPVΔ012 indicated that transcription of both *IFITM1* and *Ly6E* are dependent on DNA-PKcs (Figure 7E). Further, infection of chicken macrophages with FWPVΔ020 in WT cells showed that *IFITM1* and *Ly6E* are transcription is upregulated in chicken cells during infection with FWPVΔ020, as in CEFs (Figure 7F). As such FPV020 regulates the expression of genes that are dependent on the DNA-PK-driven intracellular signalling pathway and, as such, have the same function in FWPV as they do in VACV in inhibiting poxvirus DNA-driven innate immunity.

## Discussion

There are multiple DNA-sensing PRRs whose function contributes antiviral immunity. In general, these processes are poorly described in Sauropsida, including birds, compared to mammals, despite the ecological and economic importance of birds and their role as reservoirs of zoonotic viruses. To address this knowledge gap, we have recently described the role of cGAS/STING pathway in FWPV sensing [12] and the mechanisms of bacterial DNA sensing by TLR21 [25] in chickens. Here we extend our exploration of comparative immunology and host/poxvirus interactions by determining the contribution of DNA-PK to innate sensing of exogenous DNA and FWPV in chickens and highlight the conservation of the mechanism by which poxviruses inhibit this host response beyond the Mammalia taxon.

We found that the DNA-PK complex function in antiviral immunity is conserved between mammals and birds. Chicken cells lacking the catalytic subunit of DNA-PK, DNA-PKcs (encoded by the *PRKDC* gene) fail to produce IFN-I in response to DNA, but not RNA, and in response to poxvirus infection, equivalent to its function in mammals [10, 24]. We also show that this response requires the kinase TBK1 and the transcription factor, IRF7. This data is consistent with IRF7 being a functional replacement for IRF3 in birds and the main IRF functioning downstream of PRRs in avian antiviral immunity [25, 47]. As such, we have now demonstrated that the DNA-PK/cGAS/STING/IRF signalling pathway is functionally maintained between birds and mammals. On the other hand, other mechanisms of viral DNA sensing, for example those that use the IF16 and AIM2 inflammasome protein family [48, 49] are not used by birds as these loci are not present in avian species [50], which places to date the DNA-PK/cGAS/STING/IRF axis as the main DNA-sensing complex to be functionally available in birds.

All poxviruses produce multiple proteins that inhibit PRR-driven IFN-I production [14, 15]. This is also true of avipoxviruses, since two FWPV proteins, FWPV012 and FWPV184, are known to reduce the amount of IFN-I and ISG transcription in infected chicken cells [13, 30]. These two proteins are not thought to target DNA sensing PRRs directly, but rather to antagonise downstream signalling molecules. Here we add FPV020 to the list of FWPV interferon antagonists and identify it, along with FPV006/255, as members of the poxvirus virus OPG020/OPG031 family. The conservation of this family of poxviral immunomodulators is broader than that of OPG188a (B2/poxin) and OPG067 (E5), which target cGAMP and cGAS respectively (Figure 4A) [21, 22]. This family is not only present across multiple genera, but often has 2-3 copies per genome. VACV, for example, has two copies of OPG020 (in its ITRs) and one copy of OPG031. Similarly FPV006 and FPV006/225 are identical genes, differing in sequence from FPV020. The presence of multiple copies of these genes, along with their high degree of conservation among poxviruses, may reflect their importance in the poxvirus life cycle. The OPG020/OPG031 protein family is multifunctional. VACV OPG020 can activate the cellular hypoxic response via an interaction between its N-terminal domain and the host PHD2 protein that activates HIF-1a [44], and OPG031can inhibit NF-11B signalling via an unknown mechanism [51]. It remains to be understood whether the FPV020 and FPV006/255 encode these functions. It is noteworthy that all identified OPG020/OPG031 family proteins share the two-domain structure with an N-terminal domain that mimics mammalian prolyl-hydroxylases [44] and a C-terminal domain that has a unique fold and is responsible for interactions with Ku [17, 18]. This two-domain structure may be preserved in order to allow the dimerisation to occur that is necessary for Ku-binding, or there may be interdependent functions of the two domains that are not yet understood.

Poxviruses habitually acquire host genes by horizontal gene transfer mechanisms either directly recombination or DNA transposons, or indirectly by retrotransposons or retroviral transfer [52–55]. The copy number of these genes can be rapidly expanded, depending on the selective pressure experienced by the virus, and then the new genes evolved to provide later generations of virus with newly functional proteins [1, 53]. This method often results in protein families in poxviruses with conserved domain structures (eg BCL-2-like, Ankyrin repeat, PRANC-domain proteins) [56–58]. Here the C-terminal domains of FPV020 and FPV006/255 have the same fold (the same fold as VACV OPG020/OPG031 C-terminal domains) in part due to the LCYC motif [17, 19]. The loops and surface-exposed residues, however, are very different and likely encode the ability to bind Ku80/70, with the electronegative residues of FPV020 maintaining this interaction, but the less negatively charged loops of FPV006/255 resulting in Ku-interaction not being maintained in this protein. Targeting this conserved region of Ku is an immune evasion strategy that accommodates extensive plasticity in the OPG020/OPG031 protein family sequence, potentially allowing a large sequence space to be explored whilst retaining binding and at the same time limiting the options for host to mutate away. The success of this strategy is reflected in the conservation of the OPG020/OPG031 protein family across the poxviridae. The consistent presence of this family may also reflect a clear impact on virulence as in mouse models of VACV infection, OPG020/OPG031 knockout viruses are attenuated [51, 59].

It is estimated that the last common ancestor between avipoxviruses and orthopoxivurses split more than 200,000 years ago [60]. Combined with the consistent presence of DNA-PK inhibitors in vertebrate poxviruses, our analyses here imply that inhibition of DNA-PK is an ancient immune evasion mechanism employed by this family of viruses and supports a conserved function of DNA-PK as a positive regulator of innate immune sensing. Given the presence of viral proteins found to targeting DNA-PK in multiple dsDNA viruses [61] it is likely that this mechanism of innate immune evasion also exists across different virus families. Similar to the recent understanding of the ancient nature of cGAS signalling [62], it is clear that evolutionarily ancient mechanisms of host nucleic acid sensing are present across taxa along with viral inhibitory mechanisms that have resulted in increasingly complex mechanisms of innate sensing in mammals. As such, understanding the differences in PRRs and their variation across taxa could guide future research in wildlife immunology and significantly enhance our ability to predict the zoonotic potential of various animal hosts.

## Acknowledgements

This work was funded by UKRI grants BB/S001336/1 and BB/Y007212/1 (BF and CB), UKRI430 (BF), BB/E009956/1, BB/G018545/1, BB/H005323/1 & BB/K002465/1 (MS) and by EUROFERI (Région Centre-Val-de-Loire, France, RG).

## Supplementary Figures and Legends

**Supplementary Figure 1:**
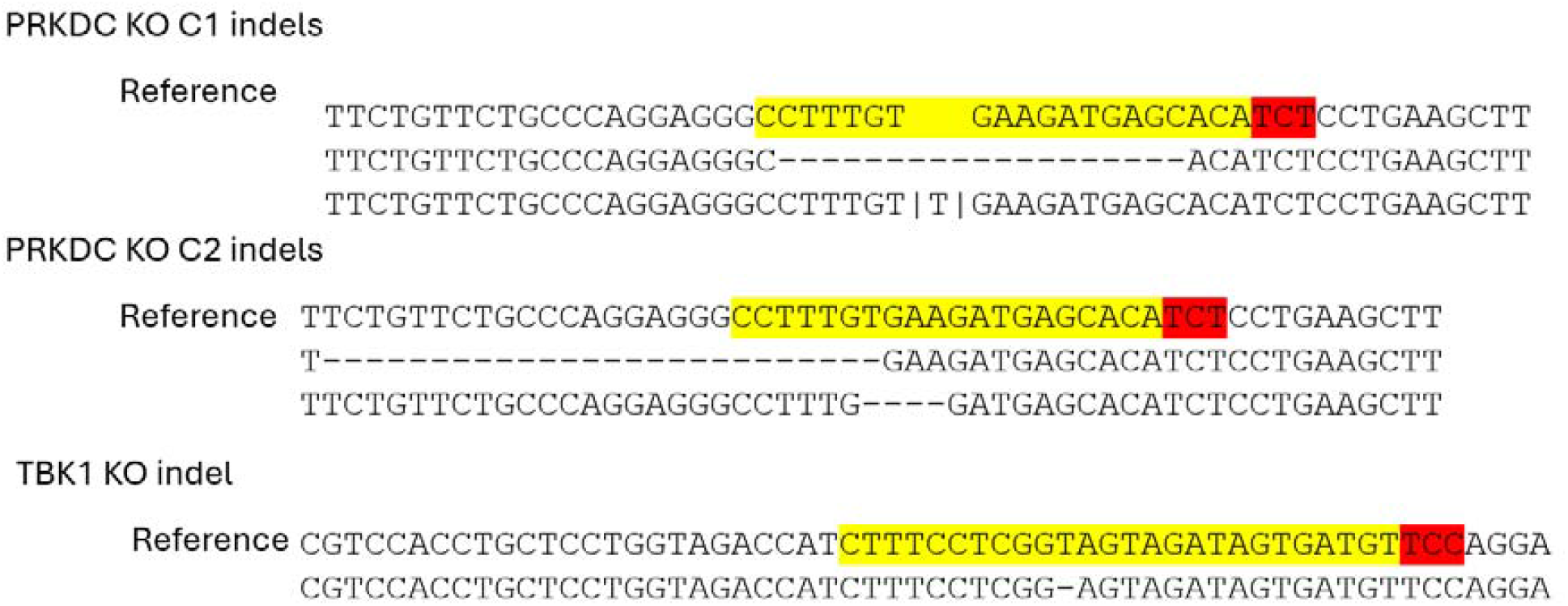
DNA-PKcs and TBK1 knockout HD11 clones were generated by CRISPR/Cas9 editing and sequenced to define indels targeted by the gRNAs (yellow) including PAM sequences (red).

**Supplementary Figure 2:**
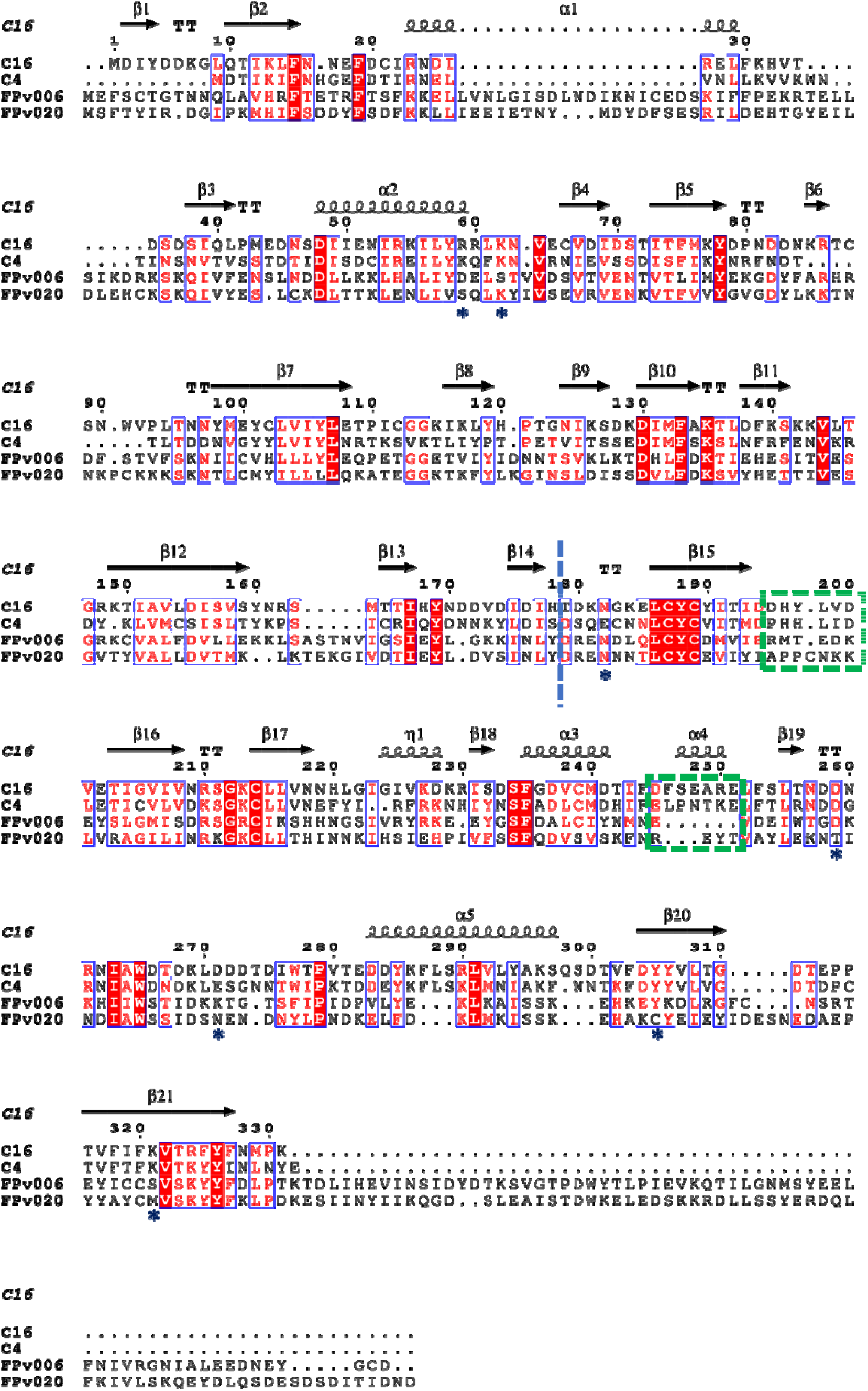
Structure based sequence alignment of VACV and FWPV C4/C10 homologues Secondary structure-guided amino-acid sequence alignments were generated based on PDB:8AG5 and AlphaFold models. Conserved residues are shown in red. Green boxes are Ku-binding loops. Blue stars indicate residues predicted to be involved in dimerisation.

## References

1. Elde NC, Child SJ, Eickbush MT, Kitzman JO, Rogers KS, et al. Poxviruses Deploy Genomic Accordions to Adapt Rapidly against Host Antiviral Defenses. Cell 2012;150:831–841.

2. Koonin EV, Dolja VV. A virocentric perspective on the evolution of life. Curr Opin Virol 2013;3:546–557.

3. Duggal NK, Emerman M. Evolutionary conflicts between viruses and restriction factors shape immunity. Nat Rev Immunol 2012;12:687–695.

4. Enard D, Cai L, Gwennap C, Petrov DA. Viruses are a dominant driver of protein adaptation in mammals. Epub ahead of print 17 May 2016. DOI: 10.7554/eLife.12469.

5. Mansur DS, Smith GL, Ferguson BJ. Intracellular sensing of viral DNA by the innate immune system. Microbes Infect 2014;16:1002–1012.

6. Bryant CE, Orr S, Ferguson B, Symmons MF, Boyle JP, et al. International Union of Basic and Clinical Pharmacology. XCVI. Pattern recognition receptors in health and disease. Pharmacol Rev 2015;67:462–504.

7. Li X-D, Wu J, Gao D, Wang H, Sun L, et al. Pivotal roles of cGAS-cGAMP signaling in antiviral defense and immune adjuvant effects. Science 2013;341:1390–1394.

8. Lam E, Stein S, Falck-Pedersen E. Adenovirus detection by the cGAS/STING/TBK1 DNA sensing cascade. J Virol 2014;88:974–981.

9. Georgana I, Sumner RP, Towers GJ, Maluquer de Motes C. Virulent Poxviruses Inhibit DNA Sensing by Preventing STING Activation. J Virol;92. Epub ahead of print 15 May 2018. DOI: 10.1128/JVI.02145-17.

10. Hristova DB, Oliveira M, Wagner E, Melcher A, Harrington KJ, et al. DNA-PKcs is required for cGAS/STING-dependent viral DNA sensing in human cells. iScience 2024;27:108760.

11. Dai P, Wang W, Cao H, Avogadri F, Dai L, et al. Modified vaccinia virus Ankara triggers type I IFN production in murine conventional dendritic cells via a cGAS/STING-mediated cytosolic DNA-sensing pathway. PLoS Pathog 2014;10:e1003989.

12. Oliveira M, Rodrigues DR, Guillory V, Kut E, Giotis ES, et al. Chicken cGAS Senses Fowlpox Virus Infection and Regulates Macrophage Effector Functions. Front Immunol 2020;11:613079.

13. Giotis ES, Laidlaw SM, Bidgood SR, Albrecht D, Burden JJ, et al. Modulation of Early Host Innate Immune Response by an Avipox Vaccine Virus’ Lateral Body Protein. Biomedicines;8. Epub ahead of print 19 December 2020. DOI: 10.3390/biomedicines8120634.

14. Smith GL, Benfield CTO, de Motes CM, Mazzon M, Ember SWJ, et al. Vaccinia virus immune evasion: mechanisms, virulence and immunogenicity. Journal of General Virology 2013;94:2367–2392.

15. Seet BT, Johnston JB, Brunetti CR, Barrett JW, Everett H, et al. Poxviruses and immune evasion. Annu Rev Immunol 2003;21:377–423.

16. El-Jesr M, Teir M, de Motes CM. Vaccinia Virus Activation and Antagonism of Cytosolic DNA Sensing. Frontiers in Immunology;11. Epub ahead of print 2020. DOI: 10.3389/fimmu.2020.568412.

17. Rivera-Calzada A, Arribas-Bosacoma R, Ruiz-Ramos A, Escudero-Bravo P, Boskovic J, et al. Structural basis for the inactivation of cytosolic DNA sensing by the vaccinia virus. Nat Commun 2022;13:7062.

18. Peters NE, Ferguson BJ, Mazzon M, Fahy AS, Krysztofinska E, et al. A mechanism for the inhibition of DNA-PK-mediated DNA sensing by a virus. PLoS Pathog 2013;9:e1003649.

19. Scutts SR, Ember SW, Ren H, Ye C, Lovejoy CA, et al. DNA-PK Is Targeted by Multiple Vaccinia Virus Proteins to Inhibit DNA Sensing. Cell Rep 2018;25:1953-1965.e4.

20. Hernáez B, Alonso G, Georgana I, El-Jesr M, Martín R, et al. Viral cGAMP nuclease reveals the essential role of DNA sensing in protection against acute lethal virus infection. Sci Adv;6. Epub ahead of print September 2020. DOI: 10.1126/sciadv.abb4565.

21. Eaglesham JB, Pan Y, Kupper TS, Kranzusch PJ. Viral and metazoan poxins are cGAMP-specific nucleases that restrict cGAS-STING signalling. Nature 2019;566:259–263.

22. Yang N, Wang Y, Dai P, Li T, Zierhut C, et al. Vaccinia E5 is a major inhibitor of the DNA sensor cGAS. Nat Commun 2023;14:2898.

23. Tao X, Song J, Song Y, Zhang Y, Yang J et al. Ku proteins promote DNA binding and condensation of cyclic GMP-AMP synthase. Cell Rep 2022;40:111310.

24. Ferguson BJ, Mansur DS, Peters NE, Ren H, Smith GL. DNA-PK is a DNA sensor for IRF-3-dependent innate immunity. Epub ahead of print 18 December 2012. DOI: 10.7554/eLife.00047.

25. Guabiraba R, Rodrigues DR, Manna PT, Chollot M, Saint-Martin V, et al. Mechanisms of type I interferon production by chicken TLR21. Dev Comp Immunol 2024;151:105093.

26. Waterhouse AM, Procter JB, Martin DMA, Clamp M, Barton GJ. Jalview Version 2--a multiple sequence alignment editor and analysis workbench. Bioinformatics 2009;25:1189–1191.

27. Katoh K, Standley DM. MAFFT multiple sequence alignment software version 7: improvements in performance and usability. Mol Biol Evol 2013;30:772–780.

28. Capella-Gutiérrez S, Silla-Martínez JM, Gabaldón T. trimAl: a tool for automated alignment trimming in large-scale phylogenetic analyses. Bioinformatics 2009;25:1972–1973.

29. Kalyaanamoorthy S, Minh BQ, Wong TKF, von Haeseler A, Jermiin LS. ModelFinder: fast model selection for accurate phylogenetic estimates. Nat Methods 2017;14:587–589.

30. Laidlaw SM, Robey R, Davies M, Giotis ES, Ross C, et al. Genetic screen of a mutant poxvirus library identifies an ankyrin repeat protein involved in blocking induction of avian type I interferon. J Virol 2013;87:5041–5052.

31. Laidlaw SM, Skinner MA. Construction of deletion-knockout mutant Fowlpox virus (FWPV). Bio Protoc;4. Epub ahead of print 20 May 2014. DOI: 10.21769/bioprotoc.1126.

32. Ge SX, Jung D, Yao R. ShinyGO: a graphical gene-set enrichment tool for animals and plants. Bioinformatics 2020;36:2628–2629.

33. Jumper J, Evans R, Pritzel A, Green T, Figurnov M, et al. Highly accurate protein structure prediction with AlphaFold. Nature 2021;596:583–589.

34. Meng EC, Goddard TD, Pettersen EF, Couch GS, Pearson ZJ, et al. UCSF ChimeraX: Tools for structure building and analysis. Protein Sci 2023;32:e4792.

35. Lees-Miller JP, Cobban A, Katsonis P, Bacolla A, Tsutakawa SE, et al. Uncovering DNA-PKcs ancient phylogeny, unique sequence motifs and insights for human disease. Prog Biophys Mol Biol 2021;163:87–108.

36. Tamura K, Adachi Y, Chiba K, Oguchi K, Takahashi H. Identification of Ku70 and Ku80 homologues in Arabidopsis thaliana: evidence for a role in the repair of DNA double-strand breaks. Plant J 2002;29:771–781.

37. Nenarokova A, Záhonová K, Krasilnikova M, Gahura O, McCulloch R, et al. Causes and Effects of Loss of Classical Nonhomologous End Joining Pathway in Parasitic Eukaryotes. MBio;10. Epub ahead of print 16 July 2019. DOI: 10.1128/mBio.01541-19.

38. Kulesza P. DNA-dependent Protein Kinase in V(D)J Recombination: Reconstitution of the Scid Defect by Human DNA-PK CDNA. 1999.

39. Sui H, Zhou M, Imamichi H, Jiao X, Sherman BT, et al. STING is an essential mediator of the Ku70-mediated production of IFN-λ1 in response to exogenous DNA. Sci Signal;10. Epub ahead of print 18 July 2017. DOI: 10.1126/scisignal.aah5054.

40. Sun X, Liu T, Zhao J, Xia H, Xie J, et al. DNA-PK deficiency potentiates cGAS-mediated antiviral innate immunity. Nat Commun;11. Epub ahead of print 1 December 2020. DOI: 10.1038/S41467-020-19941-0.

41. Abe T, Ishiai M, Hosono Y, Yoshimura A, Tada S, et al. KU70/80, DNA-PKcs, and Artemis are essential for the rapid induction of apoptosis after massive DSB formation. Cell Signal 2008;20:1978–1985.

42. Senkevich TG, Yutin N, Wolf YI, Koonin EV, Moss B. Ancient gene capture and recent gene loss shape the evolution of Orthopoxvirus-host interaction genes. MBio 2021;12:e0149521.

43. Eaglesham JB, McCarty KL, Kranzusch PJ. Structures of diverse poxin cGAMP nucleases reveal a widespread role for cGAS-STING evasion in host-pathogen conflict. Elife;9. Epub ahead of print 16 November 2020. DOI: 10.7554/eLife.59753.

44. Mazzon M, Peters NE, Loenarz C, Krysztofinska EM, Ember SWJ, et al. A mechanism for induction of a hypoxic response by vaccinia virus. Proc Natl Acad Sci U S A 2013;110:12444– 12449.

45. Giotis ES, Skinner MA. Spotlight on avian pathology: fowlpox virus. Avian Pathol 2019;48:87– 90.

46. Giotis ES, Robey RC, Skinner NG, Tomlinson CD, Goodbourn S, et al. Chicken interferome: avian interferon-stimulated genes identified by microarray and RNA-seq of primary chick embryo fibroblasts treated with a chicken type I interferon (IFN-α). Vet Res 2016;47:75.

47. Cheng Y, Zhu W, Ding C, Niu Q, Wang H, et al. IRF7 Is Involved in Both STING and MAVS Mediating IFN-β Signaling in IRF3-Lacking Chickens. J Immunol 2019;203:1930–1942.

48. Unterholzner L, Keating SE, Baran M, Horan KA, Jensen SB, et al. IFI16 is an innate immune sensor for intracellular DNA. Nat Immunol 2010;11:997–1004.

49. Hornung V, Ablasser A, Charrel-Dennis M, Bauernfeind F, Horvath G, et al. AIM2 recognizes cytosolic dsDNA and forms a caspase-1-activating inflammasome with ASC. Nature 2009;458:514–518.

50. Cridland JA, Curley EZ, Wykes MN, Schroder K, Sweet MJ, et al. The mammalian PYHIN gene family: phylogeny, evolution and expression. BMC Evol Biol 2012;12:140.

51. Ember SWJ, Ren H, Ferguson BJ, Smith GL. Vaccinia virus protein C4 inhibits NF-κB activation and promotes virus virulence. J Gen Virol 2012;93:2098–2108.

52. Bratke KA, McLysaght A. Identification of multiple independent horizontal gene transfers into poxviruses using a comparative genomics approach. BMC Evol Biol 2008;8:67.

53. Brennan G, Stoian AMM, Yu H, Rahman MJ, Banerjee S, et al. Molecular Mechanisms of Poxvirus Evolution. MBio 2023;14:e0152622.

54. Hughes AL, Friedman R. Poxvirus genome evolution by gene gain and loss. Mol Phylogenet Evol 2005;35:186–195.

55. Fixsen SM, Cone KR, Goldstein SA, Sasani TA, Quinlan AR, et al. Poxviruses capture host genes by LINE-1 retrotransposition. Elife;11. Epub ahead of print 7 September 2022. DOI: 10.7554/eLife.63332.

56. Boys IN, Johnson AG, Quinlan MR, Kranzusch PJ, Elde NC. Structural homology screens reveal host-derived poxvirus protein families impacting inflammasome activity. Cell Rep 2023;42:112878.

57. Mutz P, Resch W, Faure G, Senkevich TG, Koonin EV, et al. Exaptation of Inactivated Host Enzymes for Structural Roles in Orthopoxviruses and Novel Folds of Virus Proteins Revealed by Protein Structure Modeling. MBio 2023;14:e0040823.

58. Graham SC, Bahar MW, Cooray S, Chen RA-J, Whalen DM, et al. Vaccinia virus proteins A52 and B14 Share a Bcl-2-like fold but have evolved to inhibit NF-kappaB rather than apoptosis. PLoS Pathog 2008;4:e1000128.

59. Fahy AS, Clark RH, Glyde EF, Smith GL. Vaccinia virus protein C16 acts intracellularly to modulate the host response and promote virulence. J Gen Virol 2008;89:2377–2387.

60. Babkin IV, Babkina IN. Molecular dating in the evolution of vertebrate poxviruses. Intervirology 2011;54:253–260.

61. Hristova DB, Lauer KB, Ferguson BJ. Viral interactions with non-homologous end-joining: a game of hide-and-seek. J Gen Virol 2020;101:1133–1144.

62. Li Y, Slavik KS, Toyoda HC, Morehouse BR, de Oliveira Mann CC et al. cGLRs are a diverse family of pattern recognition receptors in innate immunity. Cell 2023;186:3261-3276.e20.

